# Ultraconserved non-coding DNA within insect phyla

**DOI:** 10.1101/696005

**Authors:** Thomas Brody, Amar Yavatkar, Alexander Kuzin, Ward F. Odenwald

## Abstract

Presence of ultra-conserved sequence elements in vertebrate enhancers suggest that transcription factor regulatory interactions are shared across phylogenetically diverse species. To date evidence for similarly conserved elements among evolutionarily distant insects such as flies, mosquitos, ants and bees, has been elusive. This study has taken advantage of the availability of the assembled genomic sequence of these insects to explore the presence of ultraconserved sequence elements in these phylogenetic groups. To investigate the integrity of fly regulatory sequences across ~100 million years of evolutionary divergence from the fruitfly *Drosophila melanogaster*, we compared *Drosophila* non-coding sequences to those of *Ceratitis capitata*, the Mediterranean fruit fly and *Musca domestica*, the domestic housefly. Using various alignment techniques, Blastn, Clustal, Blat, EvoPrinter and Needle, we show that many of the conserved sequence blocks (CSBs) that constitute *Drosophila cis*-regulatory DNA, recognized by EvoPrinter alignment protocols, are also conserved in *Ceratitis* and *Musca*. We term the sequence elements shared among these species ultraconserved CSBs (uCSBs). The position of the uCSBs with respect to flanking genes is also conserved. The results suggest that CSBs represent the point of interaction of multiple trans-regulators whose functions and interactions are conserved across divergent genera. Blastn alignments also detect putative *cis*-regulatory sequences shared among evolutionarily distant mosquitos *Anopheles gambiae* and *Culex pipiens* and *Aedes aegypti.* We have also identified conserved sequences shared among bee species. Side by side comparison of bee and ant EvoPrints identify uCSBs shared between the two taxa, as well as more poorly conserved CSBs in either one or the other taxon but not in both. Analysis of uCSBs in dipterans, mosquitos and bees will lead to a greater understanding of their evolutionary origin and the function of their conserved sequences.

## Introduction

Phylogenetic footprinting of *Drosophila* genomic DNA has revealed that *cis*-regulatory enhancers can be distinguished from other essential gene regions based on their characteristic pattern of conserved sequences (Kuzin et al. 2009; Kuzin et al. 2012) (Odenwald et al. 2005;Pennacchio et al. 2006; Brody et al. 2007; Loots and Ovcharenko, 2007; Hardison, 2000, Bergman et al. 2002). These studies have shown that most enhancers are made up of clusters of conserved sequences that often are comprised of 5 to 30 or more conserved sequence blocks (CSBs). On average, *Drosophila* enhancers span ~1 kb and are flanked by non-conserved DNA of variable length.

Cross-species alignments have also identified conserved non-coding sequence elements associated with vertebrate developmental genes (Thomas et al. 2003; Bejerano et al. 2004), and sequences that are conserved among ancient and modern vertebrates (e. g., the sea lamprey and mammals). These elements conserved between disparate phyla are considered to be ‘ultraconserved elements’ (McEwen, et al. 2009; Irvine, et al. 2002). Many of these sequences act as *cis*-regulators of transcription (Pennacchio et al. 2006; Visel et al. 2009; McEwen et al. 2009; Visel et al. 2013; Dickel, 2018). Evidence from truncation studies indicates that, in the case of a mammalian Sonic Hedgehog enhancer, the ultraconserved element is not simply a clustering of transcription factor (TF) binding sites but has a structural component that is key to its activity (Lettice et al. 2014), suggesting that such highly conserved sequence blocks fit an enhanceosome model in which multiple adjacent and overlapping transcription factor docking sites act cooperatively to regulate gene expression (Panne, 2008). Previous studies have identified ultra-conserved elements in dipterans [*Drosophila* species and sepsids and mosquitos (Glazov et al. 2005; Hare et al. 2009, Sieglaff et al. 2009, Suryamohan et al, 2016)]. Comparison of consensus transcription factor binding sites, in the spider *Cupiennius salei* and the beetle *Tribolium castaneum*, have been shown to be functional in transgenic *Drosophila* (Ayyar et al. 2010).

Adjacent CSBs within *Drosophila* enhancers exhibit evolutionary conserved spacing. For example, characterization of 19 consecutive *Drosophila* enhancers spanning ~30 Kb between the *vvl* and *Prat2* genes revealed, in many instances, an evolutionarily constrained substructure between sets of enhancer CSBs (Kundu et al. 2013). Linked associations of adjacent CSBs could also be due to fixed spatial requirements for interactions of different transcriptional regulators (see for example Gao, et al. 2008, Panne, 2008).

In this study, we describe sequence conservation between the medfly *Ceratitis capitata*, the house fly *Musca domestica* genomic sequences and *Drosophila* genomic sequences. The house fly and Medfly have each diverged from *Drosophila* for ~100 and ~120 My respectively (Beverley and Wilson, 1984). Our analysis reveals that, in many cases, CSBs that are highly conserved in *Drosophila* are also conserved in *Ceratitis* and *Musca*. Similar to ultraconserved sequences in vertebrates, we consider these cross-phyla conserved sequences to be uCSBs. Additionally, the linear order of these uCSBs with respect to flanking structural genes is also maintained. However, subset of the uCSBs exhibits inverted orientation relative to the *Drosophila* sequence, suggesting that while enhancer location is conserved, their orientation relative to flanking genes is not.

For detection of conserved sequences in mosquitos, we have adapted EvoPrinter algorithms, to include 22 species of *Anopheles* plus *Culex pipens* and *Aedes aegypti*. Use of *Anopheles* species allows for the resolution of CSB clusters that resemble those of *Drosophila*. Comparison of *Anopheles* with *Culex* and *Aedes*, separated by ~150 million years of evolutionary divergence (Krzywinski et al. 2006), reveals uCSBs shared among these taxa. Although mosquitoes are considered to be Dipterans, uCSBs were conserved between mosquito species but not with flies.

In addition, we have developed EvoPrinter tools for sequence analysis of seven bee and thirteen ant species. Both ants and bees belong to the Hymenoptera order and have been separated by ~170 million years (Peters et al. 2017). Within the bees, *Megachile* and *Dufourea* are sufficiently removed from *Apis* and *Bombus* (~100 My; Peters et al. 2017, Elsik et al. 2016) that only portions of CSBs are shared between species; these can be considered to be ultraconserved sequences. uCSBs are found that are shared between ant and bee species, and these are positionally conserved with respect to their associated structural genes. Finally, we discovered ant specific and bee specific CSB clusters that are not shared between the two taxa but are interspersed between shared uCSBs.

## Methods

### Sequence curation and alignment

*Drosophila melanogaster (Dm), Apis mellifera (Am)* and *Anopheles gambiae (Ag), the fly*, bee and mosquito genomic sequences, were curated from the UCSC genome browser. BLASTn (Altschul et al. 1990) was used to identify non-coding sequences within other species not represented in the UCSC genome browser. Where possible, BLAT (Kent, 2002) and BLASTn were used in comparing the order and orientation of ultra-conserved sequences in reference species with dipteran, bee and mosquito test species. BLAT was not available for the *Culex* comparison to *Aedes*, but we found that the ‘align two sequences’ algorithm of BLAST, using the ‘Somewhat similar sequences (BLASTn)’ setting, was comparable to BLAT in sensitivity to sequence homology and was useful in this comparison. Similarly, the pairwise sequence alignment program Needle, which uses the Needleman-Wunsch algorithm (Needleman et al. 1970), aligned shorter regions of near identity that could not be seen by other methods.

### Mosquito EvoPrinter

An EvoPrint provides a single uninterrupted view, with near base-pair resolution of conserved sequences as they appear in a species of interest. A prior paper describes protocols for genome indexing, enhanced BLAT alignments and scoring of EvoPrint alignments. Readouts are comparable to those already described (Yavatkar et al. 2008).

To compare 24 *Anopheles, Aedes* and *Culex* genomes, sequences were obtained from VectorBase (https://www.vectorbase.org/genomes). The mosquito EvoPrinter consists of 20 species, including 7 species of the Gambiae subgroup and related species *A. christyi and A. epiroticus*, 5 species of the Neocellia and Myzomyia series (including *A. stephensi, A. maculates, A. calcifacies, A. funestus* and *A. minimus*), 2 species of the Neomyzomyia series (*Anopheles darius* and *Anopheles farauti*), 2 species of subgenus *Anopheles* (*A. sinensus* and *A. atroparvus*), Nyssoryhynchus and other American species, (*A. albimanus* and *A. darling*), and two species of the subfamily Culicinae (*Aedes aegypti* and *Culex quinquefaciatus*). Mosquito genomes are documented by Holt et al. 2002; Nene, et al. 2007; Reddy, et al. 2012, and Neafsey et al. 2014.

### Hymenoptera EvoPrinter

We have also formatted seven bee species, including 6 members of the family Apidae and one member of each of the Megachilidae and Halictidae families (Table 1). In addition, we have formatted 13 ant (Formicidae) species, a diverse family of social insects, for EvoPrinter analysis (Table 1). Among these are eight representative of the subfamily Myrmicinae, three representatives of the Formicinae, two of the Ponerinae, and one Dolichoderinae. For consistency, we selected a member of the Myrmicinae as input/reference sequence, and species selection was dependent on the integrity and completeness of the sequence. The ant and bee EvoPrinter consist of the following species, grouped according to their phylogenetic relationships:

**Table 1:**
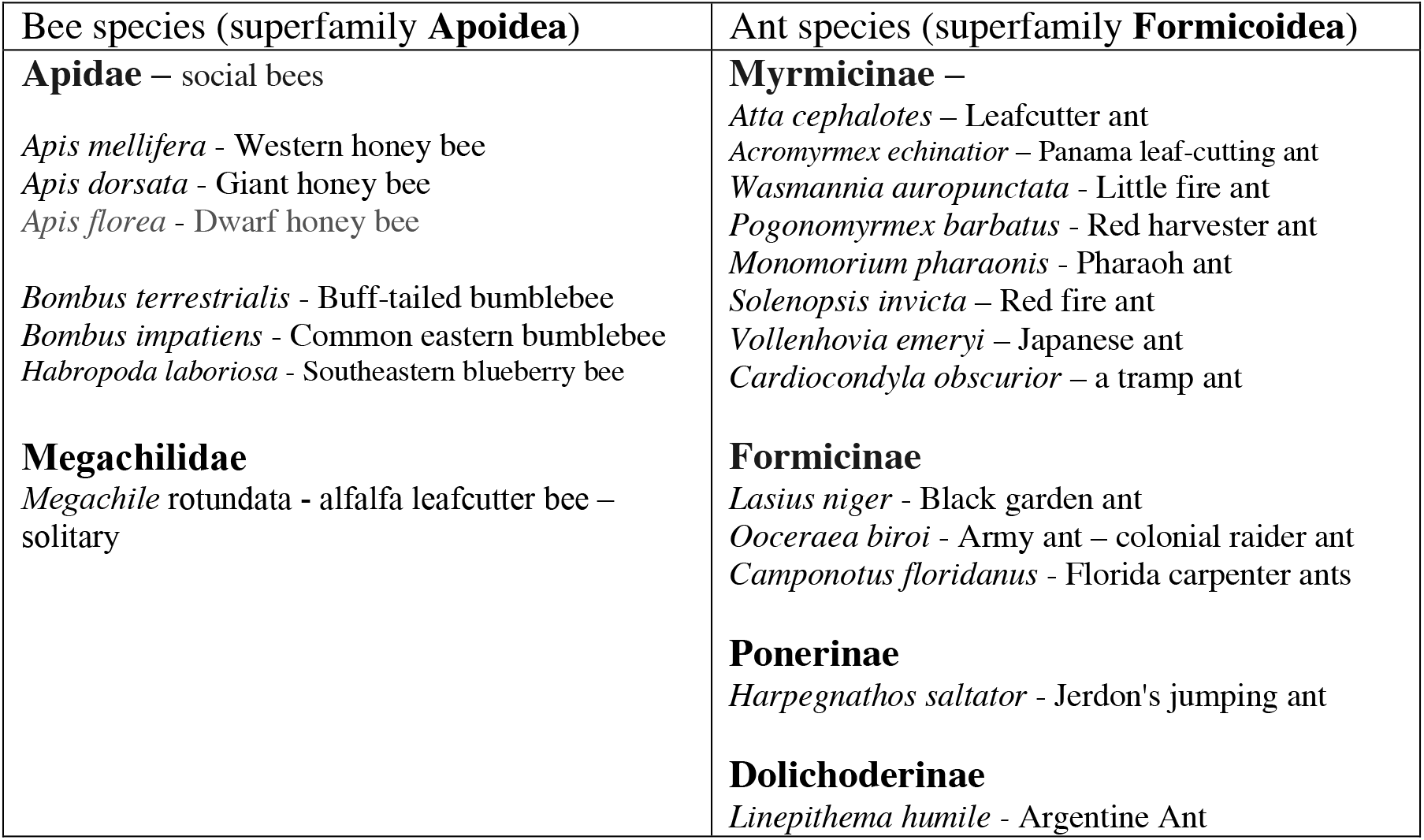
Ant and Bee species formatted for EvoPrint analysis

## Results and Discussion

### Comparative analysis of dipteran non-coding DNA

Our previous study of 19 consecutive *in vivo* tested *Drosophila* enhancers contained within a 28.9 kb intragenic region located between the *vvl* and *Prat2* genes, revealed that each CSB cluster functioned independently as spatial/temporal *cis*-regulatory enhancer (Kundu et al. 2013). The enhancers possessed a diversity of regulatory functions, including dynamic activation of expression in defined patterns within subsets of cells in discrete regions of the embryo, larvae and/or adult.

Submission of the 29 Kb enhancer field to the RefSeq Genome Database of *Ceratitis capitata* via BLASTn revealed 17 uCSBs; all 17 regions were colinear and located between the *Ceratitis* orthologs of *Drosophila vvl* and *Prat2* genes. In each case the matches between *Ceratitis* and *Drosophila* corresponded to a complete or a portion of a CSB identified as being highly conserved among *Drosophila* species (Kundu et al. 2013). Submission of the same *Drosophila* region to *Musca domestica* RefSeq Genome Database revealed 13 uCSBs that are colinearly arrayed within the *Musca* genome. Since the structural gene and these conserved uCSBs are currently on different contigs, the absolute orientation of the *Musca* sequences with respect to the *Musca vll* structural gene could not be determined. Nine of these *Ceratitis* and *Musca* CSBs were present in both species and corresponded to CSBs contained in several of the enhancers identified in our previous study of the *Drosophila* enhancer field (Kundu et al., 2013). The conservation within one of these embryonic neuroblast enhancers, vvl-41, is depicted in Fig. 1. Panel A of Fig. 1 is an EvoPrint of vvl-41 annotated to show shared CSBs with *Ceratitis* and *Musca*. Green CSBs are shared 3 ways between the three species, red letters represent bases that are shared between *Dm* and *Ceratitis* and blue letters represent bases that are shared exclusively between *Dm* and *Musca*. Fig. 1B shows two and three-way alignments in vvl-41 between the conserved CSBs in the three species. In many cases the uCSBs contained known DNA motifs for TFs. Each of the CSB elements in vvl-41 that are shared between *Dm* and *Ceratitis* are in the same orientation with respect to the *vvl* structural gene. However, in *Musca*, the orientation of elements with respect to the structural gene is unknown since the structural gene and the CSBs are on different contigs. Supplemental fig. 1 presents three-way alignments of each of the other eight uCSBs within the *vvl* enhancer field that are shared between *Dm, Ceratitis* and *Musca*. The uCSB of vvl-49 in *Ceratitis* is in reverse orientation with respect to the vvl structural gene. Many of the uCSBs in Musca are in a different orientation on the contig than in *Dm*, indicating microinversions. We conclude that, except for microinversions, the order and orientation of highly conserved non-coding sequences in *Drosophila, Ceratitis* and *Musca* with respect to flanking genes is the same.

**Figure 1.**
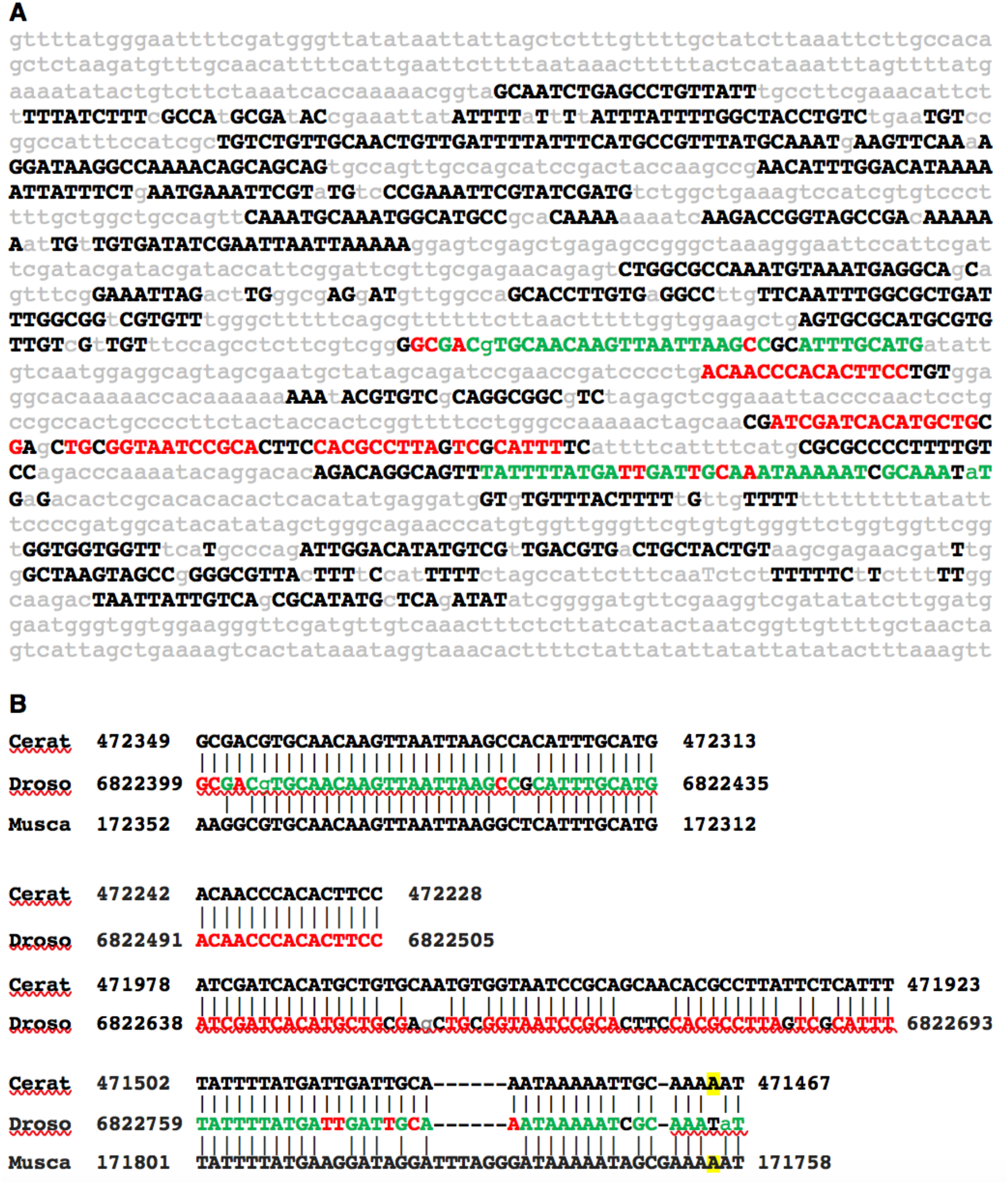
Ultra-conserved sequences shared among a *Drosophila ventral veins lacking* enhancer and orthologous DNA within the *Ceratitis capitata* and *Musca domestica* genomes. **A**) An *EvoPrint* of the *D. melanogaster vvl-41* neuroblast enhancer showing 1,775 bp, located 26.6 kb 3’ of the *vvl* transcribed sequence. Capital letters represent bases in the *D. melanogaster* reference sequence that are conserved in *D. simulans, D. sechellia, D. yakuba, D. erecta, D. ananassae, D. persimilis, D. grimshawi, D. mojavensis and D. virilis* orthologous DNAs. Lower case grey bases that are not conserved in one or more of these species. Conserved sequence blocks (CSBs) shared with *Ceratitis* and *Musca*, as detected using BLASTn, DNA Block Aligner and the *EvoPrinter* CSB aligner are shown in Green text while red bases are shared between *D. melanogaster* and *Ceratitis* but not with *Musca*. **B**) Two and three-way alignments between of the ultra-conserved CSBs using BLASTn alignments. Green and red font annotations in the *Drosophila* CSBs are as describe above. Yellow highlighted bases in *Ceratitis* and *Musca* are not shared in *Drosophila*. Flanking BLASTn designator numbers indicate genomic sequence positions.

Many of the non-coding regions in dipteran genomes contain uCSBs, especially in and around developmental determinants, and many of these are likely to be *cis*-regulatory elements such as those found in the *vvl* enhancer field. Another example is the prevalence of uCSBs found in the non-coding sequences associated the *Dm hth* gene locus. A previous study identified an ultraconserved regions in *hth* shared between *Drosophila* and *Anopheles* (Glazov et al. 2005). We have identified additional *hth* uCSBs shared among *Dm*, *Ceratitis* and *Musca*. We identified a total of 16 CSBs shared between the three species, 8 CSBs shared between *Dm* and *Ceratitis* but not *Musca*, and 7 CSBs shared between *Dm* and *Musca*, but not *Ceratitis* (fig. 2 and data not shown). Both *Ceratitis* and *Musca* contain uCSBs that were in reversed orientation with respect to the *Drosophila* orthologous regions.

**Figure 2.**
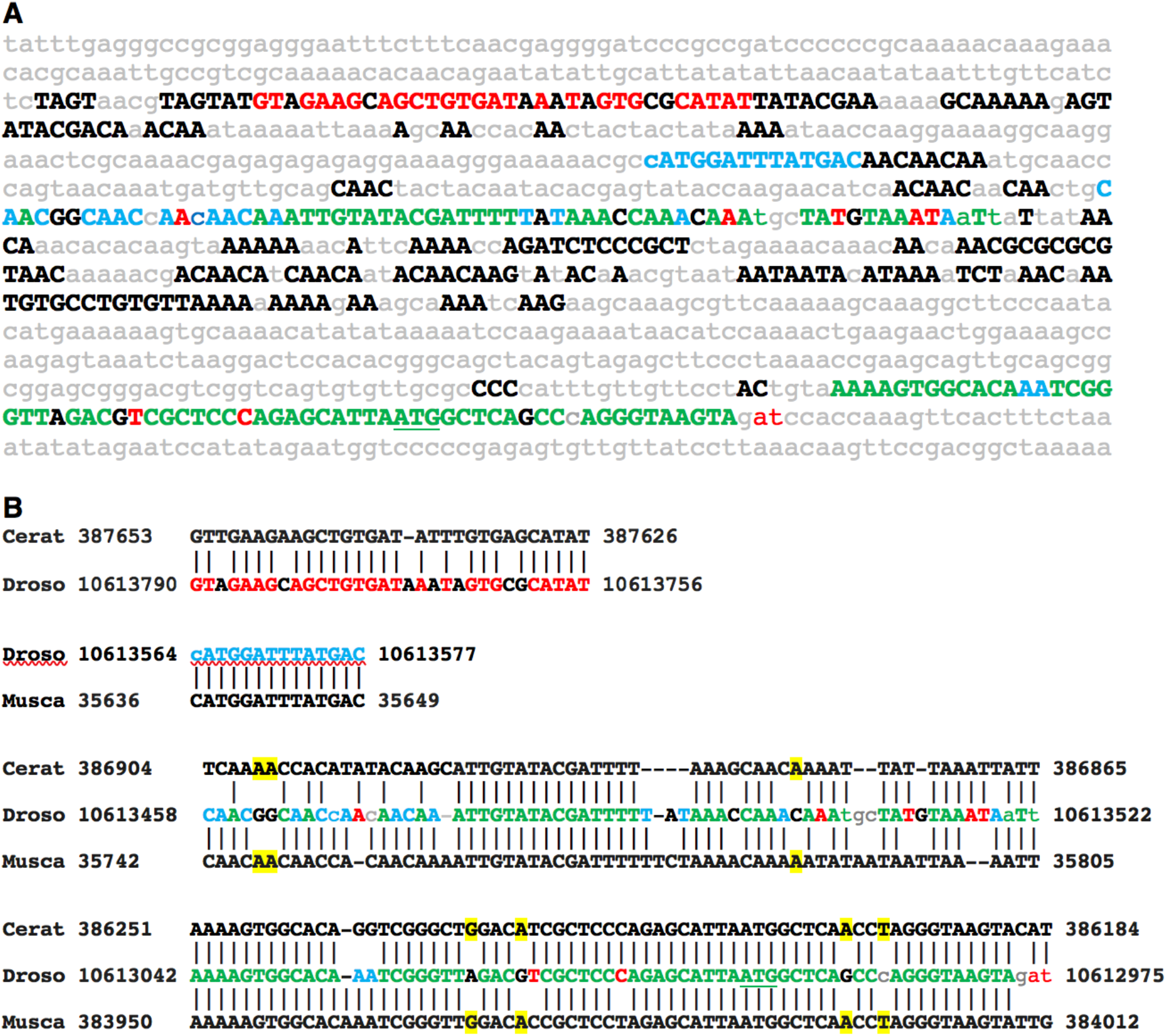
Ultra-conserved *Drosophila*, *Ceratitis capitata* and *Musca domestica* sequences within the *homothorax locus*. **A**) A 1,065bp *EvoPrint* of the *D. melanogaster homothorax locus* that includes 5’ non-transcribed sequence, its 5’ UTR, the first five codons of its encoded protein and 102bp of the first intron. Capital letters represent bases in the *D. melanogaster* reference sequence that are conserved in *D. simulans, D. sechellia, D. yakuba, D. erecta, D. ananassae, D. persimilis, D. grimshawi, D. mojavensis and D. virilis* orthologous DNAs. Lower case grey letters represent bases that are not conserved in one or more than one of the test species. *Drosophila* CSBs that are shared with *Ceratitis* and *Musca*, as detected in figure 1, are shown in green. Red bases are shared only between *Drosophila* and *Ceratitis* and blue text represent bases shared exclusively between *Drosophlia* and *Musca*. The translation start codon is marked by an underlined ATG. **B**) BLASTn two and three-way alignments of the ultra-conserved CSBs. Font color annotations are as in panel A. Yellow highlighted bases in *Ceratitis* and *Musca* are not shared in *Drosophila*. Flanking BLASTn designator numbers indicate genome base positions.

EvoPrint analysis of *Drosophila hth* sequences immediately upstream and including the first exon, revealed a conserved sequence cluster (Fig.2) associated with the transcriptional start site. Fig.2A illustrates correspondence of the *Dm* conserved region in *Ceratitis* and *Musca*. Two of the longer CSBs were conserved in both *Ceratitis* and *Musca*, one shorter CSB was conserved only in *Musca*, and a second shorter CSB was conserved only in *Ceratitis*. Two and three-way alignments as revealed by BLASTn in a comparison of *Dm*, *Ceratitis* and *Musca* are shown in Fig.2B. Each of the uCSBs is in the same orientation with respect to the *hth* structural gene.

### Discovery of non-coding conserved sequence elements in mosquitoes

EvoPrinting combinations of species using A*. gambiae* as a reference species and multiple species from the Neocellia and Myzomyia series and the Neomyzomyia provides a sufficient distance from *A. gambiae* to resolve CSBs. The CSB clusters resolved within the *Anopheles* species (data not shown) are similar to those detected using *Dm* as a reference sequence (Brody et al, 2008). Phylogenic analysis has revealed the *Anopheles* species have diverged from ~48 My to ~30 My (Kamali et al, 2014) while *Aedes* and *Culex* diversified from the *Anopheles* lineage in the Jurassic era (∼145–200 Mya; Krzywinski et al, 2006) or even earlier.

We sought to identify uCSBs in mosquitos by comparing *Anopheles* species with *Aedes* and *Culex*. We used non-coding sequences associated with the mosquito homolog of the morphogen *wingless* (reviewed by Nusse and Varmus, 1992) to discover associated conserved non-coding sequences. Fig. 3 illustrates a CSB cluster slightly more than 27,000 bp upstream of the *A. gambiae wingless* coding exons. CSB orientation in *A. gambiae* was reversed with respect to the ORF when compared to the orentations of both *Culex* and *Aedes* CSBs. We identified uCSBs, conserved in *Culex* and *Aedes*, coincide with CSBs revealed by EvoPrint analysis of *Anopheles* non-coding sequences. Supplemental fig. 2 illustrates a EvoPrinter scorecard for the non-coding *wingless*-associated CSB cluster described in Fig. 3. Scores for the first four species, all members of the gambiae complex, are similar to that of *A. gambiae* against itself, with subsequent scores reflecting increased divergence from *A. gambiae*. *Culex* and *Aedes* are distinguished from the other species by their belonging to a distinctive branch of the mosquito evolutionary tree, the Culicinae subfamily and their low scores against the *A. gambiae* input sequence. The mosquito EvoPrinter consists of 20 species, including 7 species of the Gambiae subgroup and related species *A. christyi and A. epiroticus*, 5 species of the Neocellia and Myzomyia series (including *A. stephensi, A. maculates, A. calcifacies, A. funestus* and *A. minimus*), 2 species of the Neomyzomyia series (Anopheles darius and Anopheles farauti), 2 species of subgenus Anopheles (A. sinensus and A. atroparvus), Nyssoryhynchus and other American species, (A. albimanus and A. darling), and two species of the subfamily Culicinae (Aedes aegypti and Culex quinquefaciatus). Mosquito genomes are described by Holt et al., 2002; Nene et al., 2007; Reddy et al., 2012, and Neafsey et al, 2014.

**Figure 3.**
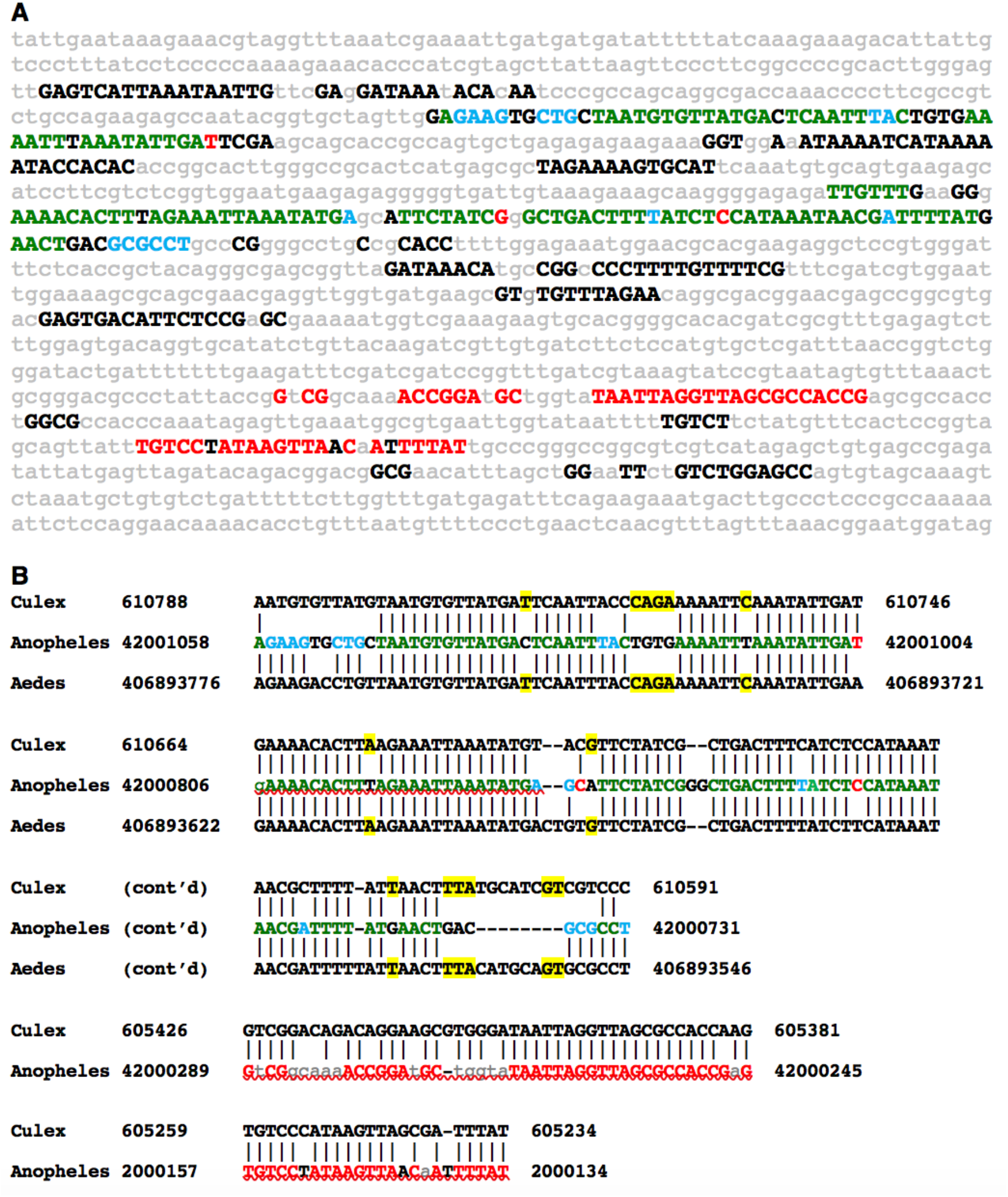
*EvoPrint* analysis of the intragenic region adjacent to the *Anopheles Wnt-4* and *wingless* genes identifies ultra-conserved sequences shared with the evolutionary distant *Culex pipiens* and *Aedes aegypti* genomes. **A**) *Anopheles gambiae* genomic *EvoPrint* that spans 1,420 bp, located 10.2 kb upstream of the *Wnt-4* gene and 27.5 kb upstream of the wingless gene which is transcribed in the opposite orientation of *Wnt-4* transcription. Capital letters (all font colors) represent bases conserved in all or all but one of the following *Anopheles* test species: *A. gambiae-S1, A. merus, A. melas, A. epiroticus, A. christyi, A. funestus, A. culicifacies, A. dirus or A. farauti.* Lower case grey letters represent bases that are not conserved in two or more of the Anopheles species included in the relaxed *EvoPrint*. Green uppercase bases indicate sequences are conserved in the *Anopheles* species, *Culex pipiens* and *Aedes aegypti*, blue font indicates *Anopheles* sequences that are shared only between *Culex pipiens* but not with *Aedes aegypti* and red font sequences are present only in *Anopheles* and *Culex*. **B**) To confirm the shared ultra-conserved CSBs, two and three-way BLASTn alignments of the shared sequences are shown. Color coding is as in panel A and yellow highlighted bases in the three-way alignments indicate identity between *Culex* and *Aedes* that is not present in *Anopheles*. Flanking BLASTn designator numbers indicate genome base positions.

### Conserved sequence elements in bees and ants

Bees and ants are members of the Hymenoptera Order, representing the Apoidea (bee) and Vespoidea (ant) super-families. Current estimates suggest that the two families have evolved separately for over 100 million years (Elsik et al. 2015: Hymenoptera Genome Database: integrating genome annotations in HymenopteraMine). To identify conserved sequences shared by bees and ants or unique to each family, we developed EvoPrinter alignment tools for seven bee and 13 ant species (Table 1). Three approaches were employed to identify/confirm conserved elements (both in coding and non-coding sequences) and their positioning within bee and ant orthologous DNAs. First, Evoprinter analysis of bee and ant genes identified conserved sequences in either bees or ants and ultra-conserved sequence elements shared by both families (figs. 4,5). Second, BLASTn alignments of the orthologous DNAs identified/confirmed CSBs that were either bee or ant specific or shared by both (data not shown). Third, side-by-side comparisons of ant and bee EvoPrints and BLASTn comparisons revealed similar positioning of orthologous CSBs relative to conserved exons (figs. 6, S2 and data not shown).

**Figure 4.**
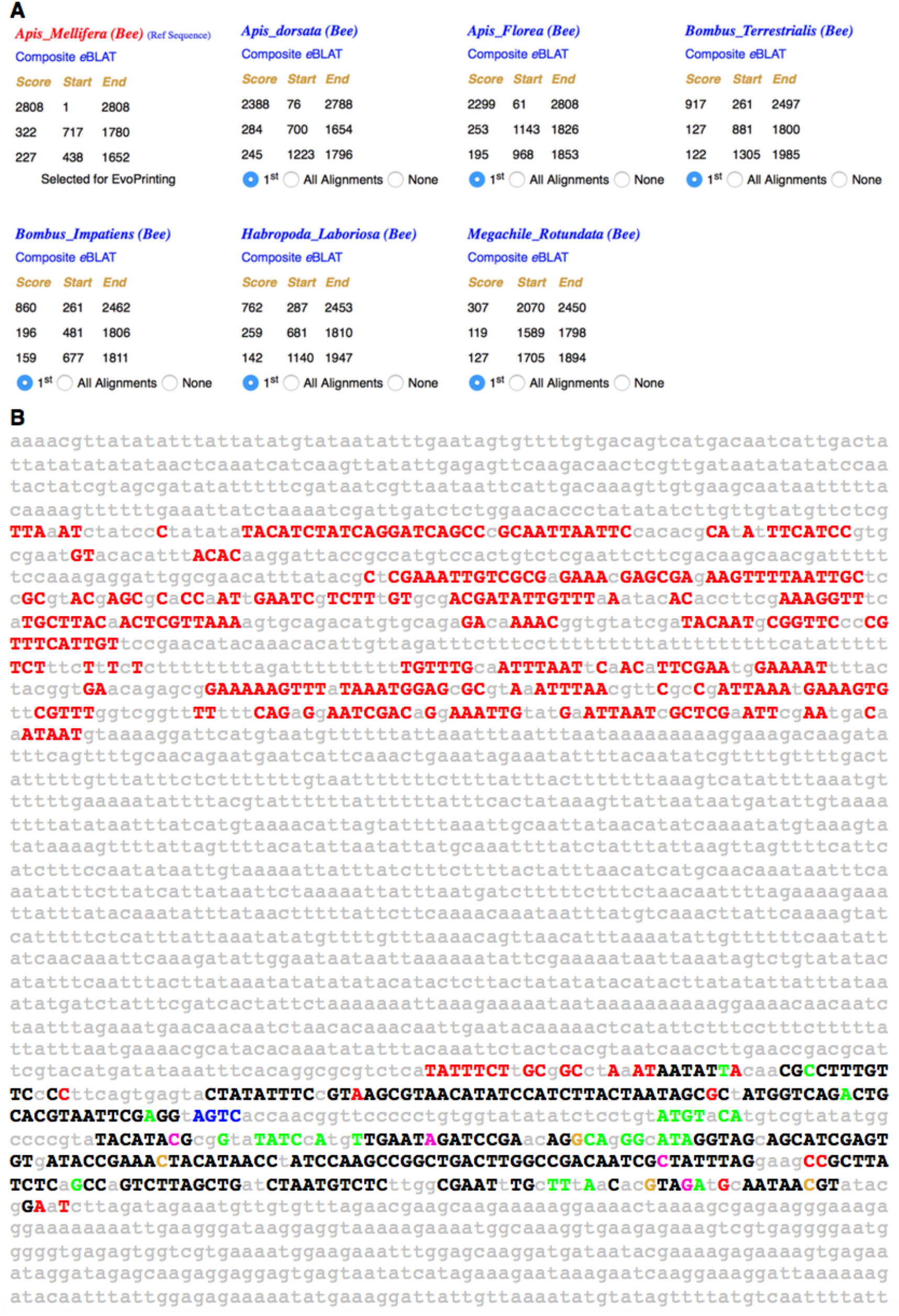
Conserved sequence clusters within the honeybee *dscam2* gene second intron. *EvoPrinter* analysis reveals *Apis mellifera* non-coding sequence elements that are conserved in other bee species or only in a subset of species. **A**) Alignment data generated from one-on-one comparisons of a 2.8 kb sequence from the honeybee 16 kb *dscam2* second intron. For each species, the top three independent *e*BLAT alignment scores are listed. Scores indicate the total number of bases within the reference sequence, the *Apis mellifera dscam2* intron, that align with the test species genome. The test species; *Apis dorsata*, *Apis florea*, *Bombus terrestrialis*, *Bombus impatiens*, *Habropoda laboriosa* and *Megachile rotundata* are listed (L -> R) based on their highest alignment score in descending order. Website links to individual *e*BLAT alignments and superimposed composite *e*BLATs are indicated in either red or blue font colors. As indicated in the alignment scorecard by the blue selection buttons, the top (highest scoring alignment) for each test species has been selected for *EvoPrinting*. **B**) A color-coded relaxed *EvoPrint* of the 2.8 kb honeybee *dscam2* second intron generated from the alignment data shown in panel A. Black uppercase letters indicate bases conserved in all test species. Font colors represent sequences conserved in all species except for *Apis dorsata, Apis florea, Bombus terrestrialis, Bombus impatiens, Habropoda laboriosa or Megachile rotundata.* Gray lowercase nucleotides are not conserved in at least two of the test species.

**Figure 5.**
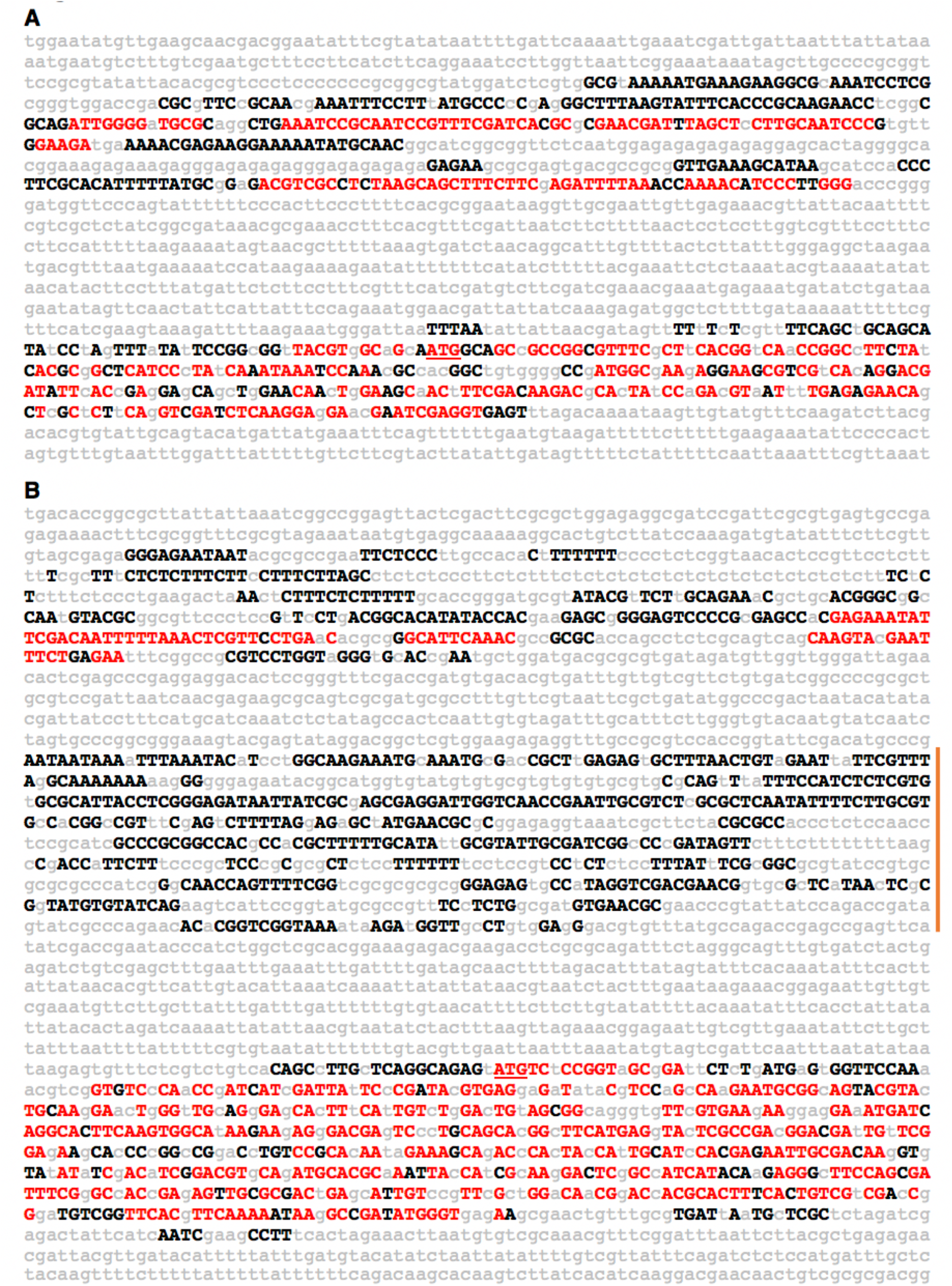
Combined Ant and Bee *EvoPrints* identify ultra-conserved *Hymenoptera* DNA. **A**) An *Apis mellifera goosecoid (gsc) EvoPrint* generated with four evolutionary divergent bee genomes and then overlaid with a print that includes the four bee genomes plus four divergent ant species. The *Apis* honeybee *gsc* DNA (1,701 bp) includes 5’ non-coding, the first exon and intron sequences. All uppercase bases (both black and red font) are conserved in bees and sequences that are conserved in both bees and ants are denoted with red-font uppercase bases. Lowercase gray-colored bases are not conserved in one or more of the bee test genomes. Bee test genomes: *Bombus terrestrialis, Bombus impatiens, Habropoda laboriosa* and *Megachile rotundata*. Ant test genomes: *Linepithema humile, Monomorium pharaonis, Wasmannia auropunctata and Atta cephalotes*. **B**) *EvoPrints* of the ant *Wasmannia auropunctata castor* (*cas*) gene locus. The 3,078 bp *Wasmannia* genomic DNA includes *cas* 5’ non-coding, the first exon and flanking intron genomic sequences. The initial *Evoprint* was generated with four evolutionary divergent ants and then super-imposed with a print that included these four ants plus four bee genomes. All uppercase bases (both black and red font) are conserved in the ants *Cerapachys biroi, Linepithema humile, Atta cephalotes and Vollenhovia emeryi*. Sequences that is conserved in both ants and bees (*Apis florea, Bombus impatiens, Habropoda laboriosa and Megachile rotundata*) are shown as red colored uppercase bases. Lowercase gray-colored bases are not conserved in one or more of the ant test species. The translation initiation codon is underlined. The left flanking vertical brown bar indicates an ant-specific conserved DNA cluster that is not found in bees. Note, in the exon ORF most, but not all, of the conserved codons do not have conserved wobble positions indicating that the cumulative evolutionary divergence of the test species used to generate the *EvoPrint* afford near base pair resolution of essential DNA.

**Figure 6.**
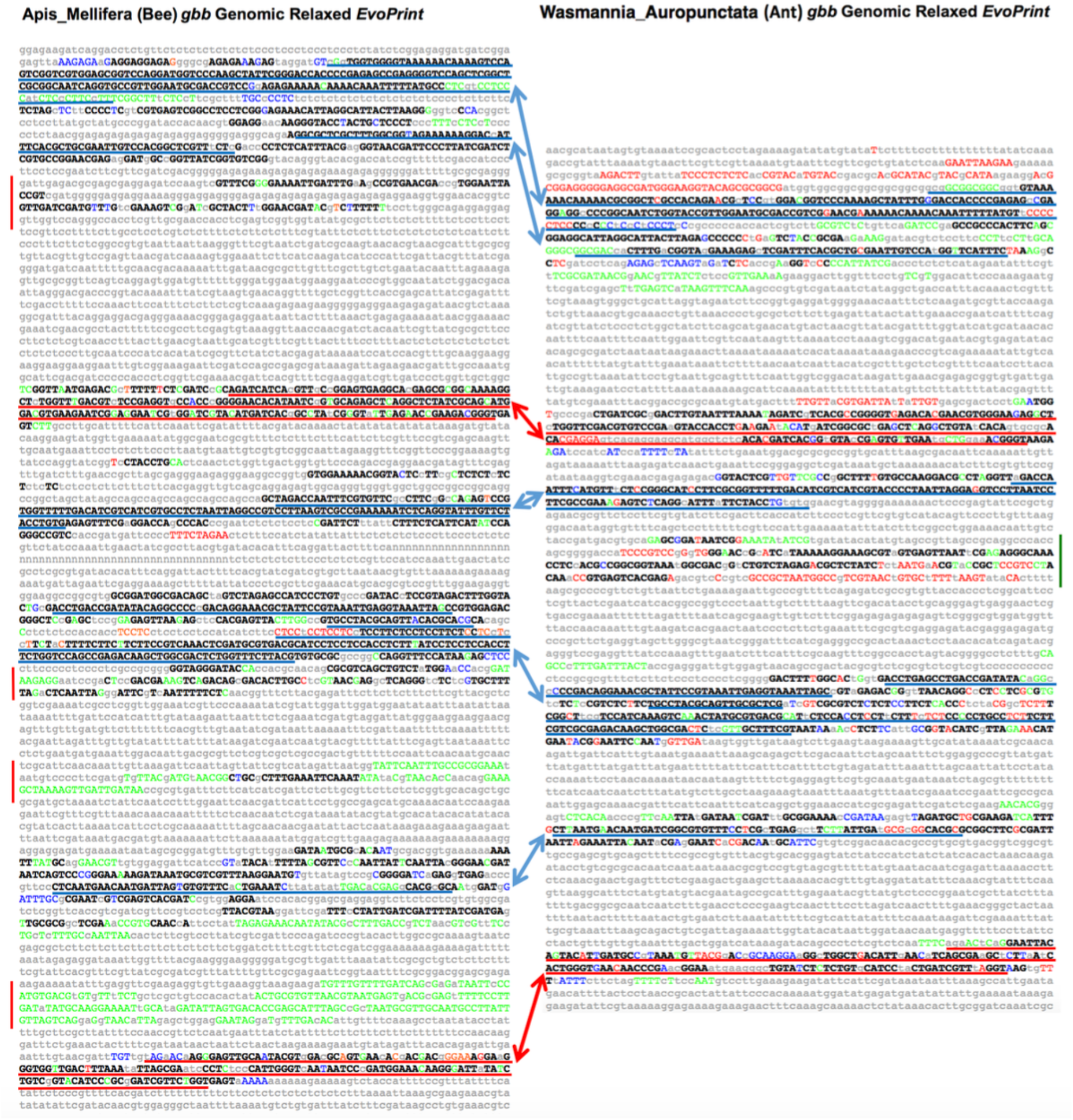
Side-by-Side comparison of conserved sequences within in the bee and ant *glass bottom boat loci* identify clusters of conserved and species-specific sequences. **A**) Relaxed *EvoPrint* of *Apis mellifera* genomic DNA that includes the *glass bottom boat* (*gbb*) second and third exons (red underlined sequences) plus flanking intronic sequences (6.6 kb). Black uppercase bases are conserved in all test bee species and colored uppercase bases are conserved in all but one of the color-coded test species: *Bombus terrestrialis, Habropoda laboriosa, Megachile rotundata* and *Bombus impatiens.* First and second exons sequences underlined red. Blue underlined sequences are homologous to underlined sequences in panel B. Vertical red bars flanking the *EvoPrint* indicate conserved bee-specific sequences that are not found in ants. **B**) Relaxed *EvoPrint* of *Wasmannia auropunctata* DNA that spans the second and third exons of the *gbb* gene including their flanking intronic sequences (5.1 kb). As in panel A, black uppercase bases are conserved in all test ant species and colored uppercase bases are conserved in all but one of the color-coded species: *Cardiocondyla obscurior, Cerapachys biroi* and *Linepithema humile.* Red and blue underlined sequences are respectively homologous coding and non-coding sequences in panel A and the green vertical bar flanking the *EvoPrint* indicates ant-specific conserved sequences that are not found in bees.

To identify conserved sequences within bee species we initially generated EvoPrints of the honey bee (Apis mellifera) genes using other *Apis* and *Bombus* species. Using EvoPrints of the *Dscam2* locus resolved clusters of conserved sequences (fig. 4). *Dscam2* is implicated in axon guidance in *Drosophila* (Millard et al. 2007) and in regulation of social immunity behavior in honeybees (reviewed by Cremer et al. 2007; Harpur et al. 2019). The EvoPrint scorecard (fig. 4A) reveals a high score (close relationship) with the homologous region in the other two *Apis* species. The more distant *Bombus* species score lower by greater than 50%, and *Habropoda* represents a step down from the more closely related *Bombus* species. *Megachile* shows a significantly lower score reflecting its more distant relationship to *Apis mellifera*. The relaxed EvoPrint readout reveals two CSB clusters (fig. 4b). Only one sequence cluster, the lower 3’ cluster, is conserved in all six test species examined, while the 5’ cluster is absent present in all species except *Megachile*. BLAST searches confirmed that the 3’ cluster was absent from *Megachile*, a more distant species *Dufourea novaeangliae*, and all ant species in the RefSeq genome database (data not shown). BLASTn alignments also revealed conservation of the 3’ cluster in the bee species *Dufourea novaeangliae*, the wasp species *Polistes canadensis* and two ant species, *Vollenhavia emeryi* and *Dinoponera quadriceps*.

*EvoPrinter* analysis of bee and ant genes that are orthologs of the *Drosophila* neural development genes *goosecoid* (*gsc*) and *castor* (*cas*) revealed conserved non-coding DNA that is unique to either bees or ants or conserved in both (fig. 5). The *Drosophila* Gsc homeodomain transcription factor is required for proper axon wiring during embryonic CNS development and has recently been linked to social immunity behavior in honeybees (reviewed by Cremer et al. 2007; Harpur et al. 2019). The *Drosophila* Cas Zn-finger transcription factor has been shown to be essential for neuroblast temporal identity decisions during neural lineage development (Baumgardt et al. 2014; reviewed by Brody and Odenwald 2007). EvoPrints of the Hymenoptera orthologs identify non-coding conserved sequence clusters that contained core uCSBs shared by both ant and bee superfamilies, and these uCSBs are frequently flanked by family-specific conserved clusters (figs. 4, 5, 6 and data not shown). For example, analysis of the non-coding sequence upstream of the *Wasmannia auropunctata* (ant) *cas* first exon identifies both a conserved sequence cluster that contains ant and bee uCSBs and an ant specific conserved cluster that has no counterpart found in bees (Fig. 5B and data not shown). It is likely that the ant specific cluster was deleted in bees, since BLASTn searchs of *Wasmannia* against the European paper wasp *Polistes dominula* reveals conservation of a core sequence corresponding to this cluster (data not shown).

The combined evolutionary divergence in the *gsc* and *cas* EvoPrints, accomplished by the using multiple test species, reveals that many of the amino acid codon specificity positions are conserved while wobble positions in their ORFs are not. The lack of wobble conservation indicates that the combined divergence of the test species used to generate the prints afford near base pair resolution of essential DNA.

Cross-group/side-by-side bee and ant comparison of their conserved DNA was performed using bee specific and ant specific EvoPrints and by BLASTn alignments (figs. 6, S2 and data not shown). Fig. 6 highlights the conservation observed among bee and ant exons and flanking sequence of the *glass bottom boat* (*gbb*, *60A*) locus of *Apis melliflera* EvoPrinted with four bee test species (panel A) and the *Wasmannia auropunctata gbb* locus EvoPrinted with three ant species (panel B). Coding sequences are underlined red, non-coding homologous regions are underlined blue, and novel CSBs present in either ants or bees but not both are indicated by the vertical lines to the side of each EvoPrint. Similarly, EvoPrinting a single exon and flanking regions of the *Apis mellifera homothorax* locus with four bee species and generating an ant specific EvoPrint of the orthologous ant sequence of the *Ooceraea biroi homothorax* locus with ten other ant species, reveals CSBs that are conserved in both *Apis* and *Ooceraea*, as well as sequences that are restricted to one of the two Hymenopteran families (supplemental fig. 2).

## Summary

Our cross-species comparisons document shared ultraconserved sequences within three separate groups of insects, e.g., flies, mosquitos and Hymenoptera. In each case, CSB clusters were shown to consist of a core of highly conserved CSBs flanked by less well conserved regions. Our previous work in *Drosophila* has shown that most CSB clusters function autonomously as enhancers that control flanking gene expression patterns. This pattern of conservation has been documented for mammalian enhancers and suggests a common structure for cis-regulatory sequences across evolution. In many cases, the uCSBs were flanked by CSBs that were not shared across phyla. We suggest that core uCSBs perform essential cis-regulatory function(s), while flanking conserved sequences, shared only by more closely related species, serve to provide the species specificity to enhancer function. Often these enhancers control a sub-pattern of gene expression. (Perry et al., 2010, Kuzin et al., 2012, Ross et al., 2015)

In the three species groups examined in this study, flies, mosquitos, and ants and bees each have similar clusters of conserved sequences. For example, the alignment of *Apis mellifera* sequences with other *Apis* and *Bombus species*, or of Anopheles gambiae with other Anopheles species resolved clusters of conserved sequences resembling in many aspects BLAT alignment of *Drosophila* Sophophora subgroup (including *D. melanogaster, D. yakuba* and *D. persimilis*) with the *Drosophila* subgroup (including *D. virilis, D grimshawi* and *D mojavensis*). These alignments revealed regions that can be considered to be, in analogy to *Drosophila*, CSB clusters flanked by regions of non-conservation (termed inter-clustal regions) (Kuzin et al. 2009; Ross et al, 2015). Adding more distantly related species, *Ceratitis* and *Musca* for flies, *Aedes* and *Culex* for mosquitos, and *Megachile* and ants for Hymenoptera revealed ultraconserved CSBs, nested within the CSB clusters. Therefore, the general pattern of conservation is the same for all three taxa examined.

In most cases both nBLAST and the EvoPrinter algorithm, based on the eBLAT algorithm had similar sensitivities and gave comparable results, but we recommend that the two techniques should be used in conjunction with one another. The advantage of EvoPrinter is the presentation of an interspecies comparison as a single alignment, while the advantage of nBLAST is that it provides a sensitive detection of sequence homology in a one-on-one alignment. EMBOSSED Needle alignment gives an even more sensitive detection of shorter sequences and is of use once BLAT or EvoPrinter has been used to discover shared CSBs and/or CSB clusters.

Consecutive CSB clusters in distantly related species are often co-linear, in that the order of is maintained with respect to flanking genes. We have documented exceptions to this in both flies and mosquitos in which mini-inversions (rearrangements) occur. The fact that the orientation of CSB clusters with respect to the ORF suggests that such inversions can be tolerated, and that the orientation is irrelevant to their putative enhancer function. However, the co-linear ordering of non-coding CSB clusters suggests that the order of CSB clusters may be important for gene regulation.

The pattern of conservation of CSB clusters in the Hymenoptera suggests that new CSB clusters have their origin not by recombination with other cis-regulatory DNA but random mutational changes. The same is true for mosquitos, in which shared sequences between *Culex* and *Aedes* are often not found in *Anopheles*. We sought to identify ultraconserved CSBs shared among bees and mosquitos that were related to those shared by *Drosophila, Ceratitis* and *Musca*, but failed to find such sequences using conventional alignment protocols. This work provides a basis for future studies to understand unique commonalities and functional differences between taxonomic groups.

## Acknowledgments

We would like to acknowledge the editorial expertise and assistance of Judy Brody, Mihaela Serpe, Rosario Vicidomini, and Saumitra Choudhury.

---

**Supplemental Figure 1.**
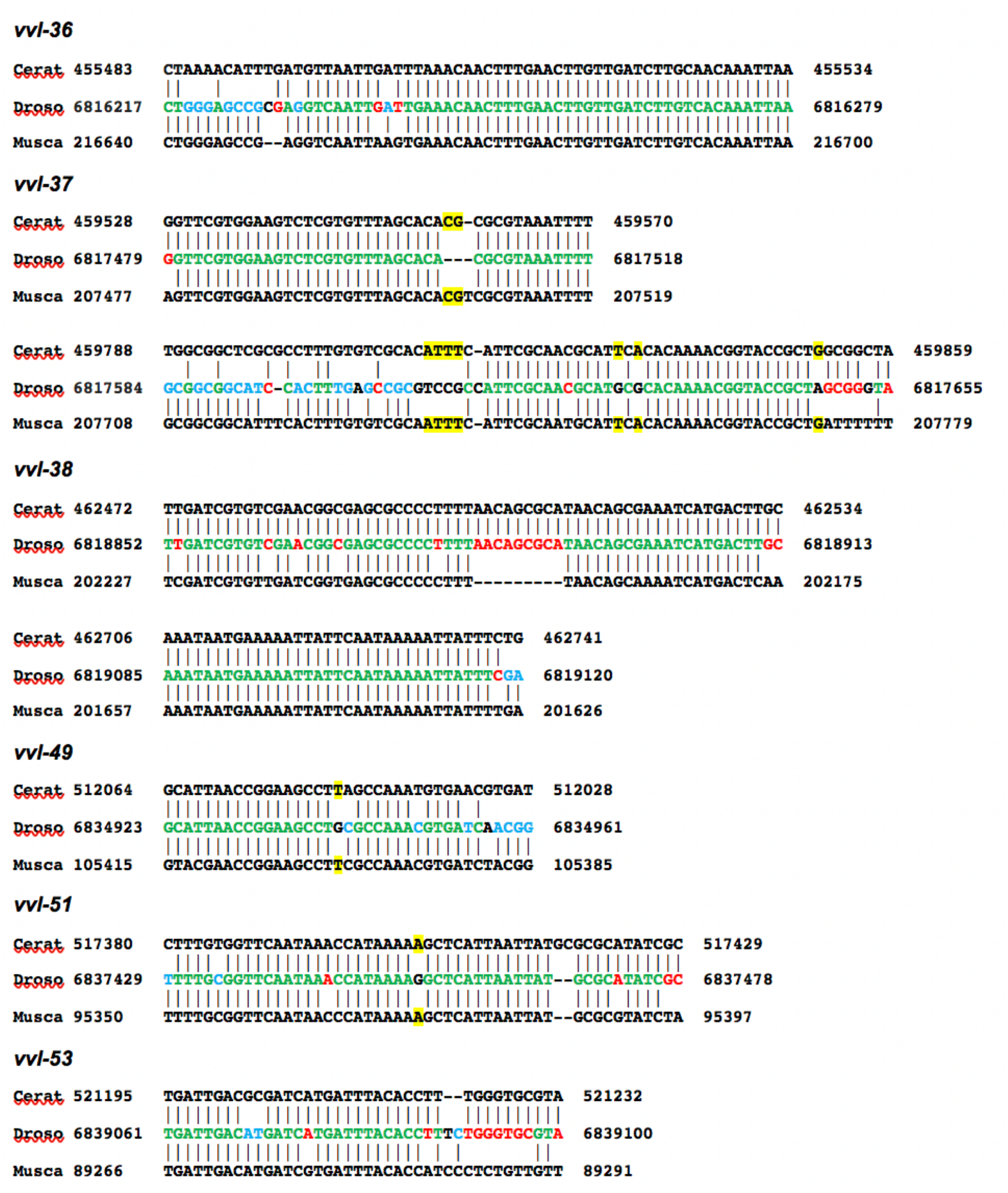
Ultra-conserved DNA in *Drosophila vvl* enhancers identified in *Ceratitis capitata* and *Musca domestica* orthologous DNAs. Three-way *Ceratitis-Drosophila-Musca* BLASTn alignments of CSBs within six different *in vivo* tested *Drosophila vvl* enhancers. *Drosophila* sequences that are shared with *Ceratitis* and *Musca* are shown in green. Red bases are shared only between *Drosophila* and *Ceratitis* and blue text represent bases shared exclusively between *Drosophila* and *Musca*. Yellow highlighted *Ceratitis* and *Musca* bases are not shared in *Drosophila*. Flanking BLASTn designator numbers indicate genomic base positions.

**Supplemental Figure 2.**
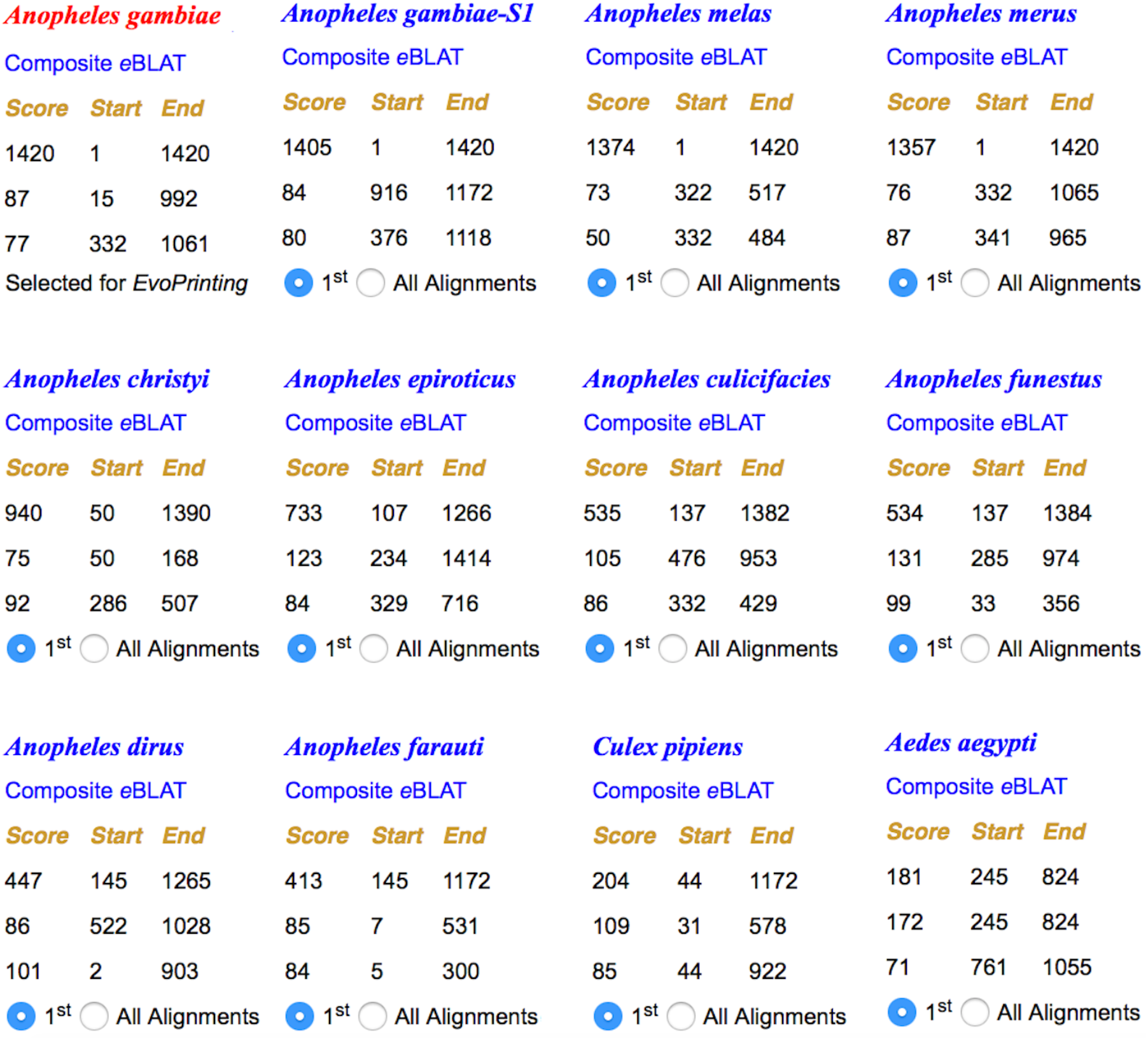
Conservation within the mosquito *wingless* gene second intron. *EvoPrinter* analysis reveals *Anopheles gambiae* non-coding sequence elements located between the mosquito homologs of Drosophila *wg* and *wnt4* that are conserved in other mosquito species. Alignment data generated from one-on-one comparisons of a 1420 base sequence from the *A. gambiae* genome. For each species, the top three independent *e*BLAT alignment scores are listed. Scores indicate the total number of bases within the reference sequence that align with the test species genome. In this analysis, 11 of the 19 mosquito test species present in the database are illustrated. The test species are listed (L -> R) based on their highest alignment score in descending order. Website links to individual *e*BLAT alignments and superimposed composite *e*BLATs are indicated in either red or blue font colors. As indicated in the alignment scorecard by the blue selection buttons, the top (highest scoring alignment) for each test species has been selected for *EvoPrinting*.

**Supplemental Figure 3.**
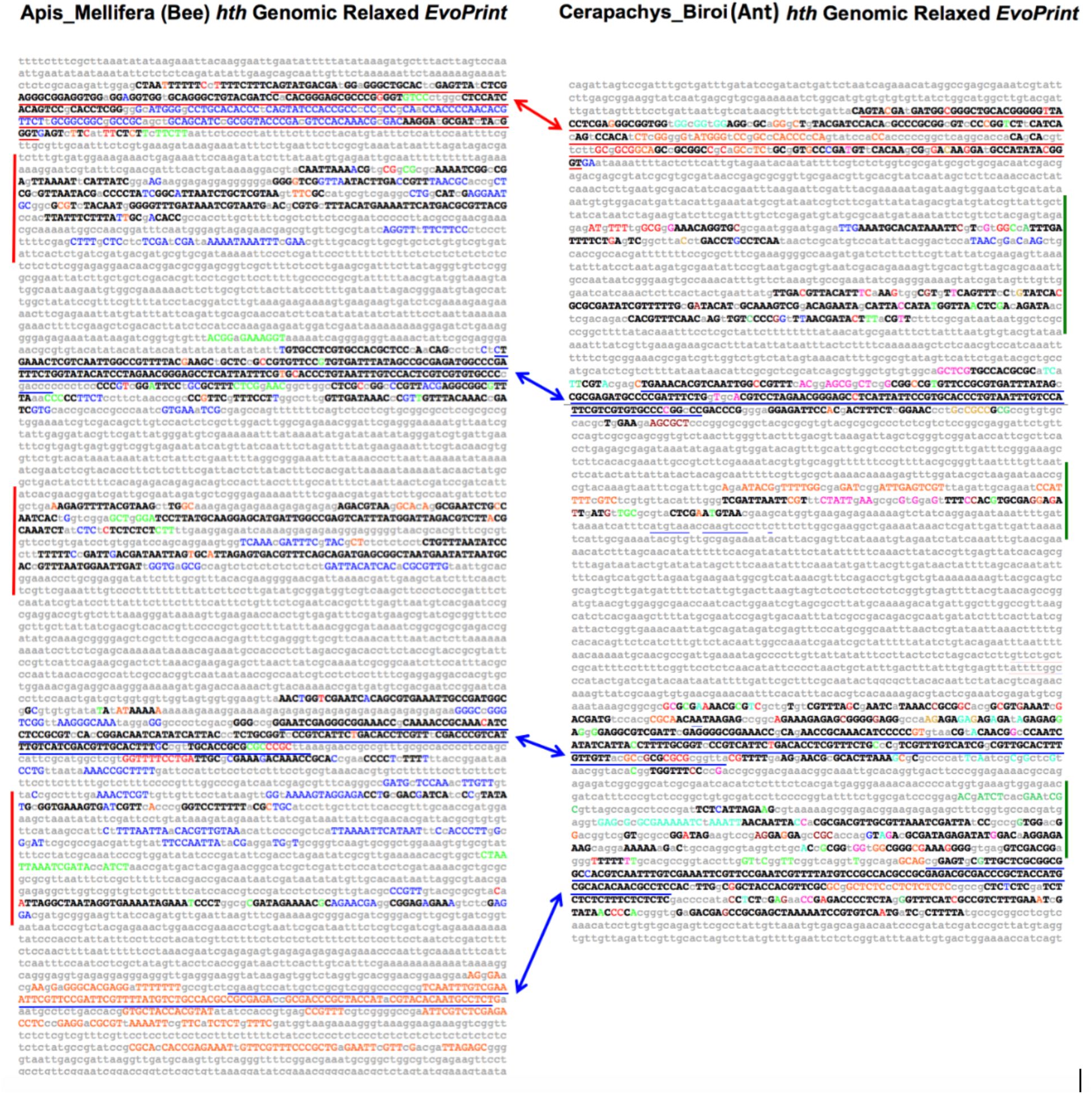
Side-by-side comparison of conserved sequences within ant and bee *homothorax* loci identifies shared exon/intron architecture and species-specific conserved sequences. *EvoPrints* of bee and ant genomic DNA that includes *homothorax* (*hth)* encoding an exon isologous to the 2^nd^ exon of *Drosophila hth* plus flanking intronic sequences. Blue and red underlined regions are coding and non-coding sequences, respectively, and align with homologous regions in the two panels. Black uppercase bases are conserved in all test species and colored uppercase bases are conserved in all but one of four bee tests species in panel A and all but one of three ant test species in panel B. **A)** Relaxed *EvoPrint* of *Apis mellifera* genomic sequences (6.3kb; Group5:7,111,526-7,117,900). Vertical red bars flanking the *EvoPrint* indicate conserved bee-specific sequences that are not found in ants. Colored uppercase bases are conserved in all but one of the color-coded test species: *Apis florea, Habropoda laboriosa, Bombus terrestrialis* and *Bombus impatiens.* **B**) Relaxed *EvoPrint* of *Cerapachys biroi* genomic DNA (5.1kb; 6532628-6527517, *Ooceraea biroi* isolate clonal line C1 chromosome 14, Obir_v5.4). The green vertical bar flanking the *EvoPrint* indicates ant-specific conserved sequence that in absent in bees. Black uppercase bases are conserved in all test ant species and colored uppercase bases are conserved in all but one of the color-coded test species: *Monomorium pharaonis, Atta cephalotes, Vollenhovia emeryi, Acromyrmex echinatior, Lasius niger, Pogonomyrmex barbatus, Wasmannia auropunctata, Cardiocondyla obscurior or Linepithema humile*.

